# Measuring speaker-listener neural coupling with functional near infrared spectroscopy

**DOI:** 10.1101/081166

**Authors:** Yichuan Liu, Elise A. Piazza, Erez Simony, Patricia A. Shewokis, Banu Onaral, Uri Hasson, Hasan Ayaz

## Abstract

The present study investigates brain-to-brain coupling, defined as inter-subject correlations in the hemodynamic response, during natural verbal communication. We used functional near-infrared spectroscopy (fNIRS) to record brain activity of speakers telling stories and listeners comprehending audio recordings of these stories. Listeners’ brain activity was correlated with speakers’ with a delay. This between-brain correlation disappeared when verbal communication failed. We further compared the fNIRS and functional Magnetic Resonance Imaging (fMRI) recordings of listeners comprehending the same story and found a relationship between the fNIRS oxygenated-hemoglobin concentration changes and the fMRI BOLD in brain areas associated with speech comprehension. This correlation between fNIRS and fMRI was only present when data from the same story were compared between the two modalities and vanished when data from different stories were compared; this cross-modality consistency further highlights the reliability of the spatiotemporal brain activation pattern as a measure of story comprehension. Our findings suggest that fNIRS is a powerful tool for investigating brain-to-brain coupling during verbal communication. As fNIRS sensors are relatively low-cost and can even be built into wireless, portable, battery-operated systems, these results highlight the potential of broad utilization of this approach in everyday settings for augmenting communication and interaction.

## Introduction

Verbal communication involves the relaying of information between individuals through the use of sound patterns within a structure of language. For decades, neuroimaging technologies have been applied to study the neural mechanisms underlying the production and comprehension of language. Multiple brain areas have been identified to be involved with verbal communication using Positron Emission Tomography (PET) and functional Magnetic Resonance Imaging (fMRI)^1^, whereas the timing of auditory processing was studied with the aid of Electroencephalogram (EEG) and Magnetoencephalogram (MEG)^2-7^. Although important findings have been discovered using these technologies, there are two limitations in traditional neurolingustic studies. First, these studies are mostly concerned with the cognitive process of either speech production or speech comprehension and confine the analysis to be within individual brains. Verbal communication, however, is an interactive process between speaker and listener. As pointed out by Hasson and others^8^, a complete understanding of the cognitive processes involved cannot be achieved without examining and understanding the interaction of neural activity among individuals. Second, cognitive functions are traditionally studied in a controlled laboratory environment. While this practice helps to isolate various factors (e.g. syntactical transformations or the representation of isolated lexical items), the ecological validity of the findings is not clear until tested in a real-life context. In addition, many studies confined the auditory stimuli to short lengths, often using isolated words or sentences for experimental control^1,7,9^. As a result, questions regarding the brain’s ability to accumulate information over longer time scales cannot be effectively investigated ^8,10,11^.

With the recent advances in neuroimaging systems and methodology, researchers can now address which brain processes are involved in social interaction. Stephens *et al.* investigated the alignment (correlation) of neural activity between speaker and listener during natural verbal communication using fMRI^12^. In the study, brain activity was recorded when a speaker was telling a real-life story and later when listeners were listening to the audio recording of the story. Listeners’ brain activity was found to be coupled with speaker’s brain activity with a delay, although for certain brain areas, listeners were ahead of the speaker in time, possibly due to a predictive anticipatory effect. Remarkably, higher coupling was found to be associated with better understanding of the story. This neural coupling between speaker and listener was further supported by a recent EEG study in which the coordination between the brain activity of speakers and listeners was investigated with canonical correlation analysis^13^. Additional findings in the same EEG study suggest that this speaker-listener neural coupling might not be restricted to homologous brain areas. In another example using fMRI, Lerner *et al.* recorded the Blood Oxygenation Level Dependent (BOLD) response of participants listening to a real-life story scrambled at the time scales of words, sentences and paragraphs. Inter-subject correlation analyses were employed to estimate the reliability of neural responses across subjects, and striking topography differences in brain activation were found at the different time scales^14^.

Although novel findings have been discovered using fMRI and EEG to address the aforementioned challenges, certain limitations of the two neuroimaging technologies have hindered the investigation of neural coupling during natural verbal communication. fMRI, for example, requires subjects to lie down motionlessly in a noisy scanning environment. Simultaneous scanning of multiple individuals, engaged in a face-to-face communication are impractical for fMRI based setups. EEG, on the other hand, was able to provide a more naturalistic environment. However, EEG is susceptible to muscle induced artifacts during vocalization, and is therefore less suitable for studying speaker-listener interactions. Furthermore, the source of the EEG signal cannot be reliably localized despite substantial efforts in the community to solve the inverse problem^15,16^.

In this study, we propose using functional near-infrared spectroscopy (fNIRS) to investigate speaker-listener coupling as an effective complement to the existing studies. fNIRS is an optical brain imaging technology for monitoring the concentration changes of oxygenated hemoglobin (ΔHbO) and deoxygenated hemoglobin (ΔHbR) in the cortex. By utilizing portable, safe and wearable sensors, fNIRS provides a cost-effective and easy-to-use imaging solution for studying brain activation in real-life contexts^17^. fNIRS has been adopted to study brain-to-brain coupling during a cooperation-competition game^18^ and a finger-tapping imitation task^19^. fNIRS has demonstrated usefulness for studying social interactions in a natural setting^20^. At present, studying brain-to-brain coupling during natural verbal communication using fNIRS has not been demonstrated.

The main objective of this study is to evaluate the feasibility of fNIRS as a new tool to study speaker-listener neural coupling. To achieve this objective, we designed an fNIRS experiment to replicate the speaker-listener neural coupling results from a previous fMRI study^12^. An English speaker and two Turkish speakers told an unrehearsed real-life story in their native language. An additional real-life English story E2 (“Pie-man”, recorded at “The Moth”, a live storytelling event in New York City) used in recent fMRI studies of natural verbal communication^14,21^, was also used here. The resulting two English stories (E1 and E2) and two Turkish stories (T1 and T2, the control conditions) were played to listeners who only understand English. We hypothesized that: 1) Neural activities of the listeners demonstrate inter-subject correlation during comprehension of only the same English stories; 2) Neural activities of the English speaker during production of E1 are coupled with the activities of the listeners during comprehension; and 3) fNIRS biomarkers (ΔHbO and ΔHbR) (recorded in this study) are correlated with the fMRI BOLD response (recorded in the previous fMRI study^14^) during the comprehension of only the same English story E2.

## Results

### Listener-listener fNIRS inter-subject correlation

For each story, significantly coupled optodes were identified using multilevel general linear model (GLM) and the results are shown in Figure 1. As expected, significant results were found only for the English conditions E1 and E2 (false discovery rate^22^ [FDR] _q_ <0.01), indicating that neural coupling only emerges during successful communication (i.e. when subjects understand the story content). ΔHbO shows a much stronger coupling effect than ΔHbR.

**Figure 1.**
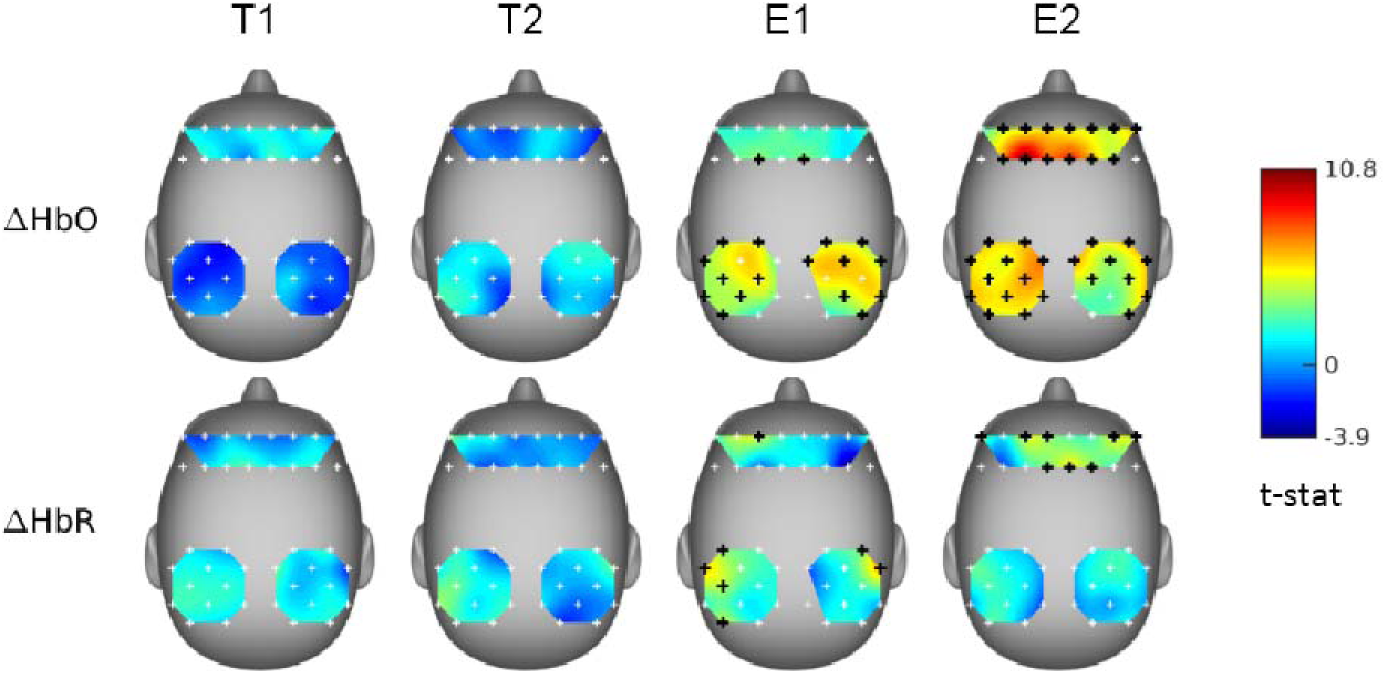
*Listener-listener fNIRS inter-subject correlation. The top row shows the t-Map of coupling results from ΔHbO and the bottom row shows the results from ΔHbR. White crosses represent non-significantly coupled optodes; Black crosses represent significantly coupled optodes (n=15, FDR q* < 0.01). *The t-Maps were smoothed using a spline method*.

### Speaker-listener fNIRS coupling

The neural coupling between speaker and listener may not be restricted to homologous brain areas^13^. Previous studies also showed that the neural responses of listeners can lag behind^12,13^ or precede^12^ those of the speaker, facilitating comprehension and anticipation, respectively. To investigate these effects, multilevel GLM has been adopted to evaluate the coupling between all permutations of (speaker optode, listener optode) pairs with the speaker’s time course shifted with respect to those of the listeners from −20s to 20s in 0.5s increments, where a positive shift represents the speaker preceding (listener lagging), and the results are shown in Figure 2. For the English story E1, the listeners’ fNIRS signals were found to be significantly coupled with the speaker’s signal with a 5-7s time delay, and the number of significantly coupled optodes peaked at 5s (Figure 2a). As expected, no temporal asymmetry has been found for the listener-listener case and alignment is coupled to the incoming auditory input (i.e. lag 0, moment of vocalization) (Figure 2b). The speaker-listener lagged correlation replicated the speaker-listener lagged correlated observed with fMRI^12^. Significant couplings can mainly be found between prefrontal of speaker and parietal of listeners in the medial prefrontal and left parietal areas for ΔHbO and no results was significant at FDR *q* < 0.01 level for ΔHbR (Figure 2c). No significant speaker-listener coupling was found for either of the Turkish stories (stories T1 and T2).

**Figure 2.**
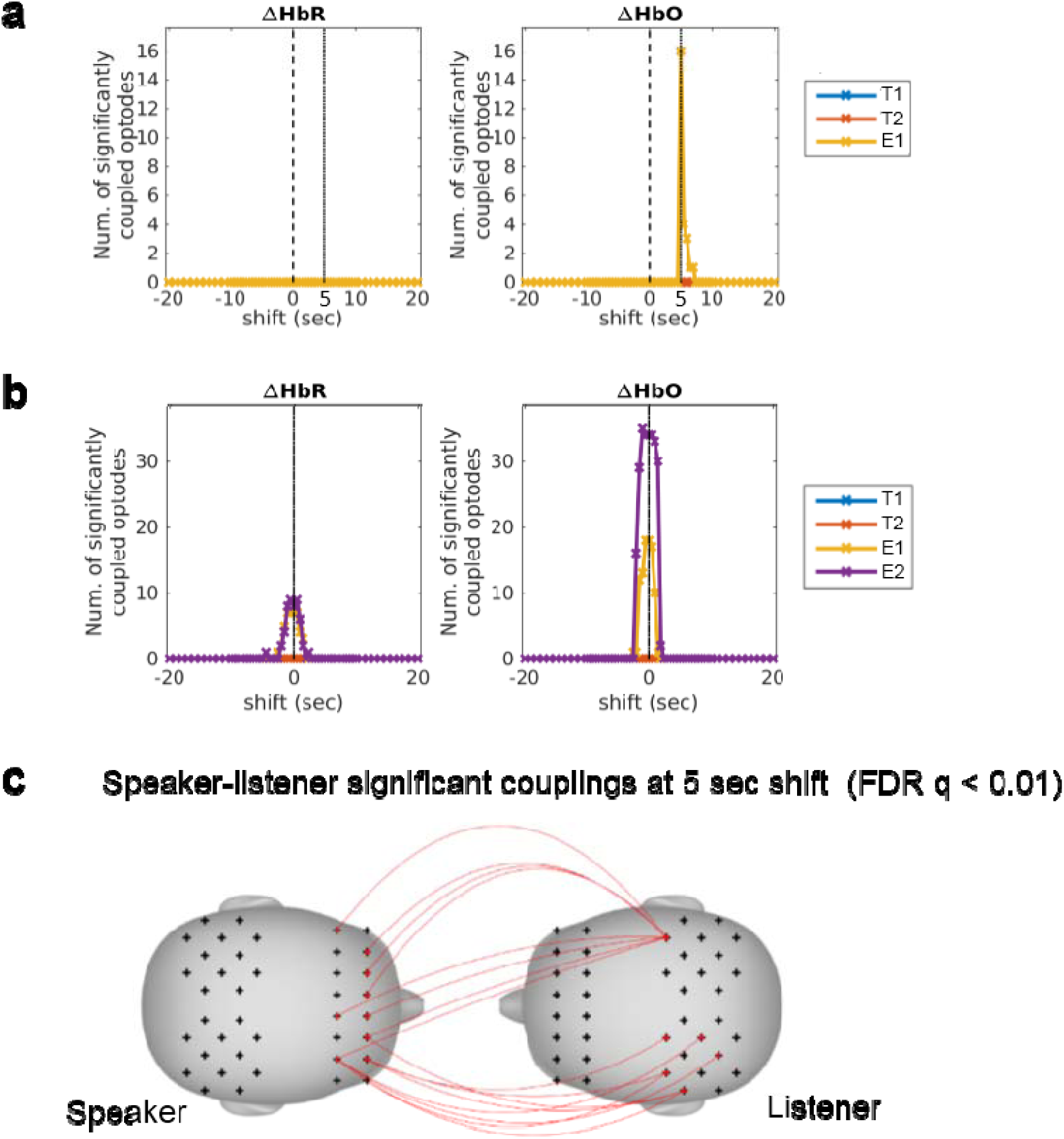
*Speaker-listener neural coupling. **a:** The number of optode-pairs at which the fNIRS time courses between a speaker and the listeners were significantly coupled (n=15, FDR _q_* < 0.01). *Results are shown with the speaker’s time course shifted from −20 to 20 sec. with respect to the listeners’ in 0.5 s increments. Significant results can be found from 5 to 7 sec. of shift (speaker precedes) with a peak at 5 sec for the English condition E1. **b:** The number of optodes showed significant listener-listener inter-subject correlation (n=15, FDR _q_* < 0.01). *Results are shown with the average listener’s time course shifted from −20 to 20 sec. with respect to those of each individual listener in 0.5s increments. No temporal asymmetry effect can be found. **c:** Significantly coupled speaker-listener optode-pairs were in non-homologous areas (n=15, FDR q* < 0.01).

To further investigate the temporal asymmetry of coupling, the average t-statistics for all significantly coupled speaker-listener optode-pairs were assessed across time shifts between the speaker and listener time series, as shown in Figure 3 (red curve). The peak of the curve is centered at 5 seconds, which shows that, on average, listeners’ time courses lagged behind the speaker’s. In comparison, the time courses of the listeners were synchronized (with each other) at 0 sec (Figure 3, blue curve).

**Figure 3.**
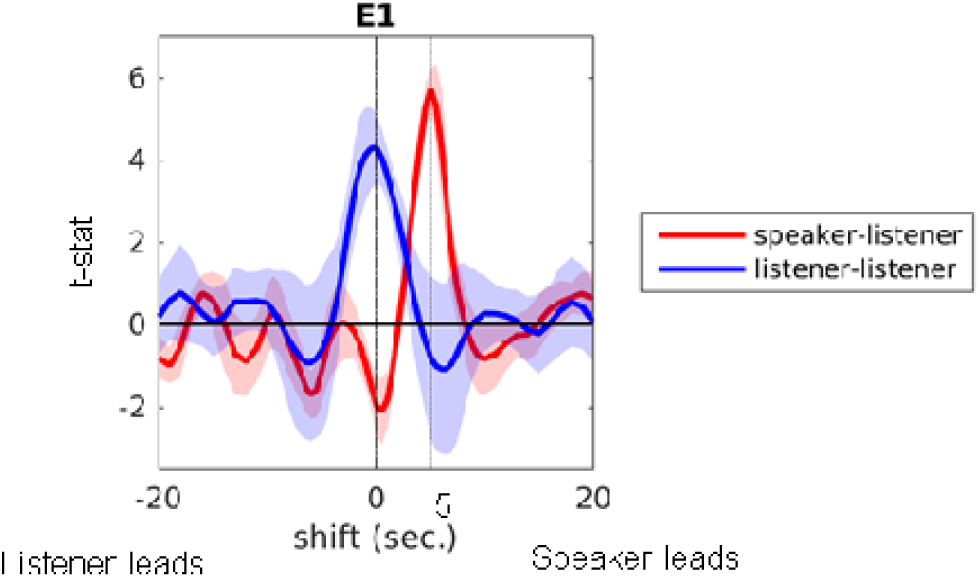
Delayed synchrony between speaker and listener. The mean distribution of t-values across significantly coupled optodes for speaker-listener (red) and listener-listener (blue) analyses. Results are shown with the speaker’s (or average listener’s) time course shifted from −20 to 20 sec. with respect to the listeners’ in 0.5 s increments. A larger t-statistic indicates stronger synchrony between the signal time courses. Listener-listener coupling was centered at 0 sec whereas speaker-listener coupling was centered at 5 sec. This suggests that, on average, the speaker preceded, and listeners needed time to process the information conveyed in the stories in order to synchronize with the speaker. Similar results have been found previously^12,13^.

### Listener-listener BOLD coupling

As a verification of the fNIRS approach, we reanalyzed an fMRI dataset of 17 subjects listening to story E2 (the e “Pie-man” story), which was recorded and used in a previous study^14^. To compare with the fNIRS results, we we considered only voxels from the outer layer of the cortex in the neighboring regions of prefrontal and parietal sites. The coupling results estimated using the multilevel GLM model are shown in Figure 4. Of the 994 investigated ted voxels, 551 showed significant listener-listener coupling (FDR. This result replicates published result^14^ and demonstrate a nice convergence across fNIRS and fMRI methods.

**Figure 4.**
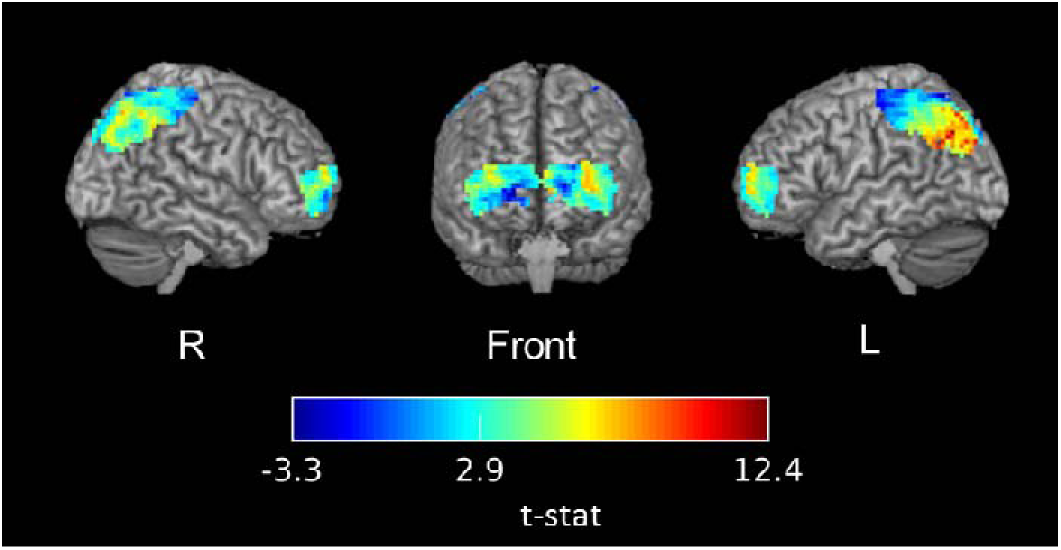
T-map of listener-listener coupling during the comprehension of story E2 (“Pie-man” story) evaluated with fMRI. The significance threshold was at t(16)=2.9 (FDR. The t-statistics were superimposed on the ch2better template and rendered using MRIcron software^42^.

### BOLD and ΔHbO are correlated during comprehension of the same story

Previous studies have shown that fNIRS and fMRI signals are highly correlated across multiple cognitive tasks^23-26^. In our study, two groups of subjects, the brain activity of one group measured with fNIRS and the other with fMRI, were engaged in the same task of listening to the E2 story (“Pie-man”). We hypothesize that the BOLD and fNIRS signals share common information even though they were measured from different subjects and in different recording environments. To directly compare the signals across fMRI and fNIRS, we estimated correlations between spatially overlapping voxel-optode pairs while subjects listened to the exact same story. Widespread significant correlations can be found between BOLD and ΔHbO only when the participants were listening to the same story (i.e. E2) as shown in Figure 5a.

**Figure 5.**
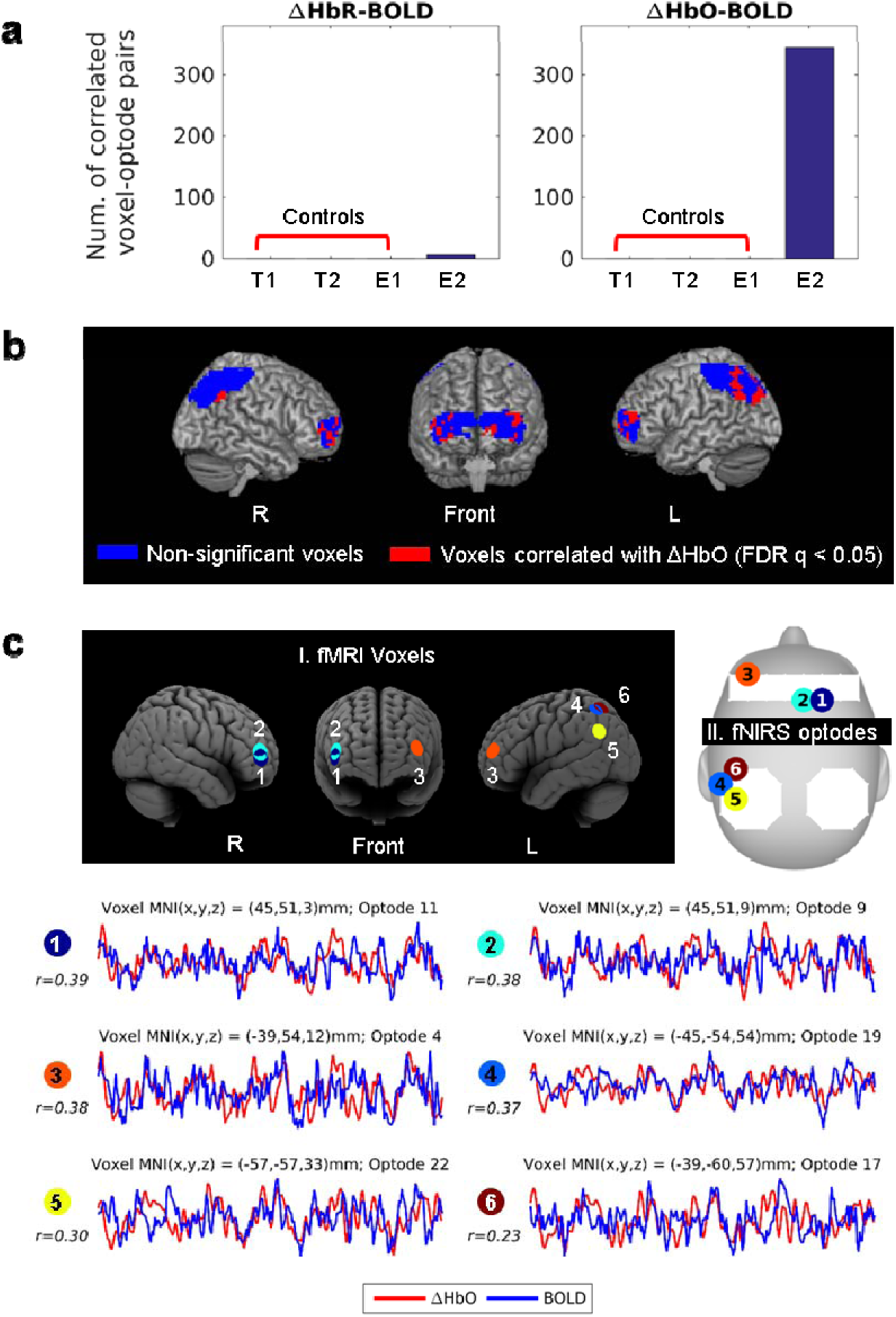
Correlation between fNIRS optodes and corresponding fMRI voxels. 994 related voxels were selected as described in section 0. **a:** The number of significant BOLD-ΔHbO and BOLD-ΔHbR correlations (n=15, FDR); here we compared the fMRI time series from story E2 with the fNIRS time series from stories E1, E2, T2, and T2. When comparing time series corresponding to different stories, we found no significant correlations between BOLD and fNIRS. **b:** Brain maps showing voxels correlated with the ΔHbO of at least one optode (red) and voxels with no significant correlations with fNIRS (blue). The images were rendered with MRIcron^42^. **c**: Six examples of voxel-optode pair. Three of the examples are the most correlated within prefrontal and the other three most correlated within parietal. Each colored circle represents one voxel-optode pair. For each pair, the voxel and optode locations are illustrated in **I. fMRI Voxels** and **II. fNIRS optodes** respectively, and the BOLD (blue line) and ΔHbO (red line) time course during the comprehension of story E2 are compared (duration = 385 s). Time courses were first standardized and averaged across the subjects and the Pearson’s correlation ( ) between BOLD and ΔHbO were estimated.

## Discussion

During social interaction, the brains of individuals become coupled as those individuals send and receive signals (light, sound, etc.) through the environment, analogous to a wireless communication system^8^. This brain-to-brain coupling relies on the stimulus-to-brain coupling which reflects the brain’s ability to be coupled with the physical world in order to represent it, veridically and dynamically. In this study, we identified the brain-to-brain coupling between a speaker telling a real-life story and a group of listeners listening to the story, with the aid of fNIRS. We also compared the brain-to-brain coupling between people listening to a real-life story across two modalities, fNIRS and fMRI. Coupling was found to be reliable using either fNIRS or fMRI, and strong correlations were found even across the two modalities. In the following sections, the implications of our findings are discussed.

### fNIRS is a viable tool for studying brain-to-brain coupling during social interaction

In this study, we demonstrated for the first time that it is feasible to study neural coupling during natural verbal communication with fNIRS. While there is a growing literature using fMRI and EEG to study brain-to-brain coupling during social interaction^27^, the application of fNIRS in the field is still rare. Cui *et al.* in 2012 first adopted fNIRS to investigate neural coupling between pairs of subjects playing a simple cooperation and competition game^18^. In the game, participants were asked to press a response key after a ‘go’ signal, either in synchrony with (cooperation mode) or faster than (competition mode) their partner. An increase in neural coupling between the members of a pair was found only during cooperation and was associated with better cooperation performance. Holper *et al.* in 2012 investigated neural coupling between a model and an imitator during a finger tapping task ^19^. A stronger increase in neural coupling was found during the imitation condition compared with a control condition in which the imitator no longer needed to follow the model’s tapping pace. These two studies, however, involved only simple stimuli and contexts. Our study demonstrated that: 1) the brain activation recorded by fNIRS was coupled between a speaker telling a real-life story and listeners listening to the story; 2) on average, the listeners’ brain activity lagged behind that of the speaker; 3) the brain activity evoked by the same story was reliable across the listeners; and 4) coupling was not present when listeners heard stories in a language incomprehensible to them. These findings are consistent with previous work using fMRI^12^ and EEG^13^ and demonstrate that fNIRS as a promising tool for studying neural coupling during social interaction in a natural communicative context.

For the speaker-listener coupling, an interesting observation is that significant inter-subject correlations were found primarily between prefrontal areas in the speaker and parietal areas in the listeners. This result supports a previous EEG study in which the coupling between speaker and listener was found to be mainly between different channels^13^ In the study, listeners watched the video playback of speakers who were telling either a fairytale or the plot of their favorite movie or book. A canonical correlation analysis between the EEG of the speakers and listeners showed that coupling was mainly limited to non-homologous channels. EEG, however, suffers from volume conduction effect. Each EEG channel records a mixture of activities from the entire brain and it is difficult to localize the correlated brain areas between speaker and listener. Comparatively, the high spatial specificity of fNIRS makes it a more suitable tool to study the non-homologous coupling phenomenon. To our best knowledge, our study presents the first fNIRS-based evidence that the brain-to-brain coupling between speaker and listener were mainly between non-homologous brain areas.

The current study, however, is only the first step toward studying neural coupling during natural communication using fNIRS. While fNIRS have obvious disadvantages relative to fMRI, which include coarser spatial resolution and an inability to measure signal beneath the cortical surface, it also has some advantages over fMRI. The advantages of fNIRS over fMRI include: 1) lower cost; 2) easier setup; 3) greater ecological validity, which can allow face-to-face communication (in fMRI setups subjects can’t see each other); 4) easier to connect two systems simultaneously to test bi-directional dialogue-based communication; and 5) easier to be used in clinical real-life communication settings. In recent studies, fNIRS has been used in extremes such as in aeurospace applications^28^ and with mobile participants walking outdoors^29^. In the context of the current study, the true advantages of fNIRS can be exploited when the neural activation of two or more subjects is studied during face-to-face conversations in a natural context such as a classroom.

Despite these promising results, our study was limited in certain aspects. First, for the fNIRS-fMRI comparison, the spatial resolution and coverage of our fNIRS system were limited, so fMRI-fNIRS correlations were estimated between all possible voxel-optode pairs within large cortical areas or the whole brain. Second, our data from fNIRS and fMRI were not recorded simultaneously and involved different participants, so any possible between-subject differences should be taken into account when interpreting the results. Future studies, preferably with concurrent fNIRS and fMRI, can validate our findings and deepen our understanding of the fNIRS-fMRI relationship for complex natural stimuli.

### fNIRS and fMRI signals are correlated for the same natural stimulus

In this study, the fMRI and fNIRS signal time courses were compared when two groups of subjects were listening to the same audio recording of a real-life story. The neural activation of one group was recorded with fNIRS. The neural activation of the other group (the fMRI group) was recorded with fMRI in a previous study^14^. We first analyzed the two datasets separately with inter-subject multilevel GLM and found similar patterns of coupling between listeners in the two modalities. We then investigated the correlation between fMRI and fNIRS signals and found that ΔHbO and BOLD were significantly correlated, despite the fact that they were collected from different subjects in different recording environments, and with different techniques (fMRI vs. fNIRS). Furthermore, the fMRI voxels that were significantly correlated with fNIRS optodes were not randomly distributed but came from brain areas usually considered to be related to listening comprehension^1^. When the fNIRS and fMRI signals corresponding to different stories were compared for control purposes, no significant correlation was found, as expected. Significant BOLD-ΔHbR correlations were found but to a much lesser extent compared to ΔHbO (Figure 5a). One possible explanation is the superior reliability of ΔHbO across the listeners compared to ΔHbR as shown in Figure 1.

For many years, researchers have been interested in comparing the fNIRS and fMRI responses during cognitive tasks. Strangman *et al.* simultaneously recorded fNIRS and fMRI in a finger flexion/extension task^30^. Although the authors expected the ΔHbR-BOLD correlation to be strongest due to the causal relation between the ΔHbR and BOLD signals, the results suggested stronger BOLD-ΔHbO correlations. The authors suspected this might be due to a higher signal-to-noise ratio (SNR) of ΔHbO in response to the task. Cui *et al.* simultaneously recorded fNIRS and fMRI for the same group of subjects during 4 tasks: finger tapping, go/no-go, judgment of line orientation and visuospatial n-back^23^. They found that both ΔHbR and ΔHbO were correlated with BOLD, despite differences in SNR, and the type of task did not significantly affect the correlations. ΔHbO-BOLD correlations were found to be slightly but significantly higher than ΔHbR-BOLD. Noah *et al.* compared fMRI and fNIRS measurements in a naturalistic task in which participants played the video game Dance Dance Revolution and rested alternately for 30-second blocks, and the results suggested a high ΔHbO-BOLD correlation within the same measurement areas ^24^. All of the studies above compared the mean triggered average activity induced by averaging a condition over time. However, Ben-Yakov *et al.* demonstrated the shortcoming of the triggered averaging method for detecting an events’ specific responses which are locked to the structure of each particular exemplar^21^. In our study, each event in the story is unique and singular and can not be averaged with the responses evoked by other events in the story. Our findings provide evidence that the response dynamics to a sequence of events in the story are robust and reliable, and can be detected both with fMRI and fNIRS, by measuring the reliability (correlation) of responses to the story within and between the two methods in real-life complex settings that unfolds over several minutes.

In summary, our results showed that: 1) A speaker’s and listener’s brain activity as measured with fNIRS were synchronized only when the listeners understood the story; 2) listening to a real-life story evoked at least some common brain activation patterns across listeners that were independent of the imaging technology (fNIRS or fMRI) and recording environment (quiet or noisy; sitting down or lying down); and 3) fNIRS and fMRI signals were correlated during the comprehension of the same real-life story. These results support fNIRS as a viable future tool to study brain-to-brain coupling during social interaction, in real-life and clinical settings.

## Materials and methods

### Participants

Three speakers (one male native English speakers, two male Turkish speakers) and 15 native English listeners (8 females) volunteered to participate in the study and were included in the analysis. An additional six subjects participated in the study but were excluded from analysis due to technical issues during recording or the excessive motion artifact presented in the large sensor array data that covers both prefrontal and parietal cortices. Subjects were all right-handed (mean LQ = 74.5, SD = 24.1) based on Edinburgh Handedness Inventory^31^ and ages 18-35 years. All subjects had normal or corrected-to-normal vision. Participants did not have any history of neurological/mental disorder and were not taking any medication known to affect alertness or brain activity. None of the listeners understood Turkish. The protocol used in the study was reviewed and approved by the Institutional Review Board (IRB) of the Drexel University (DU). The methods were carried out in accordance with approved ed guidelines and participants gave written informed consent approved by the IRB of DU.

### Experimental Procedure

All participants were seated comfortably in front of a computer screen throughout the experiment. fNIRS data of the three speakers were recorded while they told an unrehearsed real-life story in their native language (either English or Turkish). Audio of the stories was recorded using a microphone. The resulting one English story (E1) and two Turkish story (T1 and T2) were played to the listeners later. An additional real-life English story E2 (“Pie-man”, recorded at “The Moth”, a live storytelling event in New York City) used in several recent fMRI studies of natural verbal communication^14,21^, was also played to the listeners.

fNIRS data were recorded from the listeners throughout the audio playbacks. The playback sequence always ys began with E2 (English story, Pie-man), and order of the remaining stories (E1, T1, and T2) was counterbalanced ed across subjects. Before each story playback, short samples of scrambled audio were played to the subjects so they ey could adjust the volume of the headphones they were wearing. Before the start and after the end of the audio story ry playback, there was a 15-s fixation period for stabilizing the signal. Immediately after each playback, subjects were re asked to write a detailed report of the story they just heard to verify if they understood the story. Figure 6, below w, shows the timeline of a story session.

**Figure 6.**
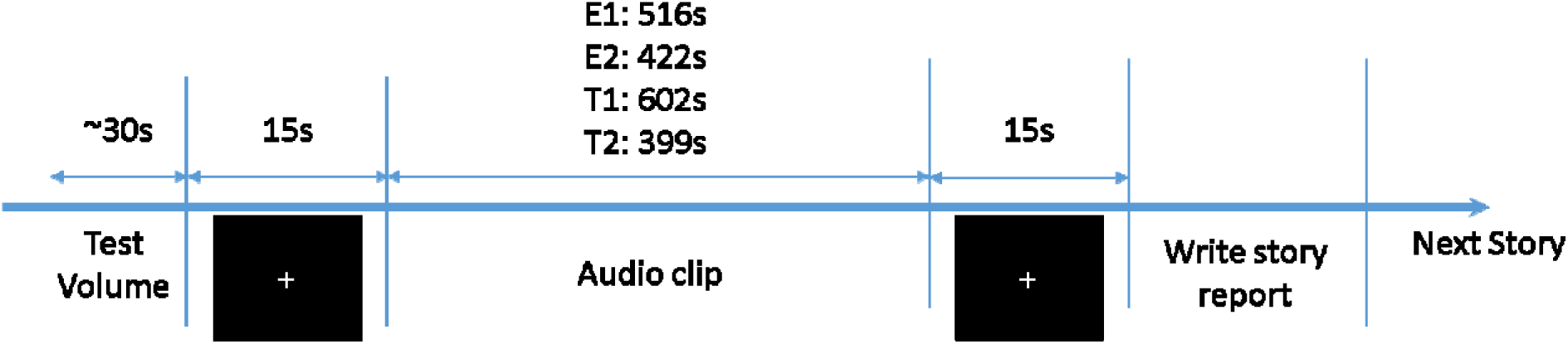
Story presentation timeline

### Data Acquisition

Two optical brain imaging devices were used simultaneously on each participant to record brain activity from prefrontal cortex (PFC) and parietal cortex (PL) using 40 measurement locations (optodes). Prefrontal and parietal regions were selected based on the significant areas found in the previous fMRI-based speaker-listener neural coupling study by Stephens, et al. ^12^. Anterior prefrontal cortex was recorded by a 16-optode continuous wave ve fNIRS system (fNIR Imager Model 1100; fNIR Devices, LLC) first described by Chance *et al.*^32^ and developed in our lab at Drexel University^33,34^. The sensor was positioned based on the anatomical landmarks as described before re in Ayaz *et al.*^34^. Briefly, the center of the sensor was aligned to the midline and the bottom of the sensor was touching the participant’s eyebrow so that the center point of the sensor was approximately at Fpz according to the 10-20 international system (see Figure 7). The sampling rate was 2 Hz. Parietal cortex was recorded using a 24-optode Hitachi fNIRS system (ETG 4000; Hitachi Medical Systems). Two “3×3” measurement patches were attached to a cap that was customized for the measurement of the parietal cortex. For each subject, the center of the two patches was placed at Pz, which was located using a measuring tape. Sensors from each patch measured the fNIRS signal of one hemisphere from 12 channels. The sampling rate was 10 Hz. Figure 7 shows the complete sensor setup and optode configuration.

**Figure 7.**
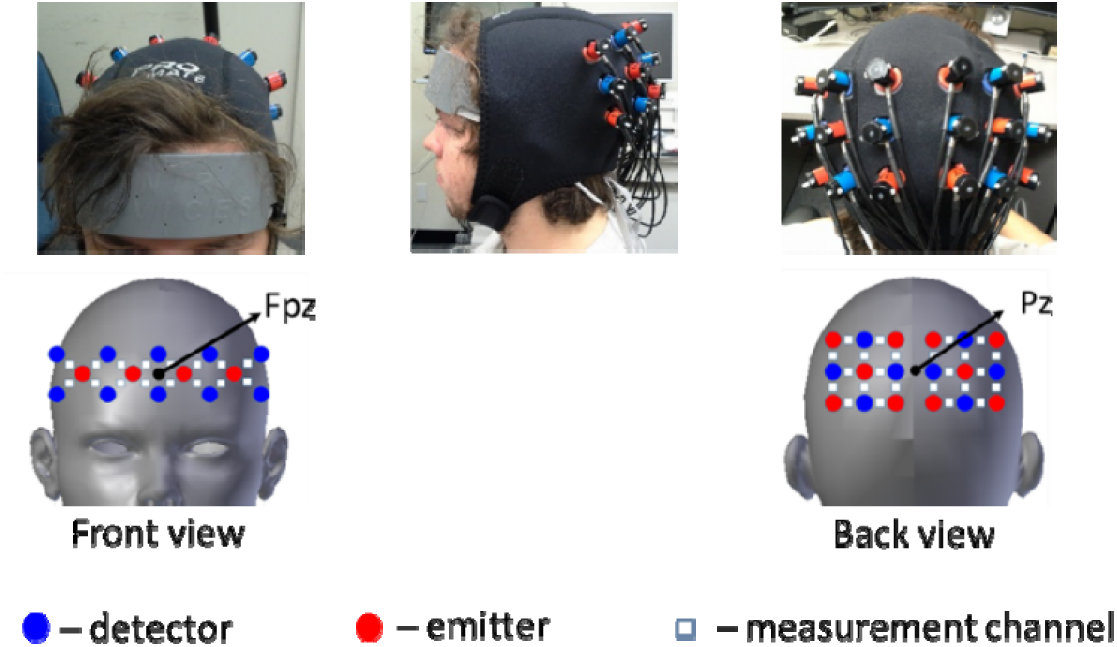
fNIRS acquisition setup. Red circles indicate emitters; blue circles indicate detectors; White squares indicate measurement channels between emitters and detectors

The approximate projection of the channel locations onto the cortical surface in MNI space was estimated using a virtual spatial registration approach^35,36^. In this approach, the sensor patches are virtually placed on an ideal scalp and the projected Montreal Neurological Institute (MNI) coordinates on the cortical surface and the standard deviation of displacement were estimated from the magnetic resonance (MR) images of 17 individuals that were obtained from a publicly available dataset^37,38^. The results are shown in Figure 8. The optodes covered regions in the e frontopolar area, orbitofrontal area, dorsolateral prefrontal cortex, primary somatosensory cortex, somatosensory association cortex, supramarginal gyrus and angular gyrus.

**Figure 8.**
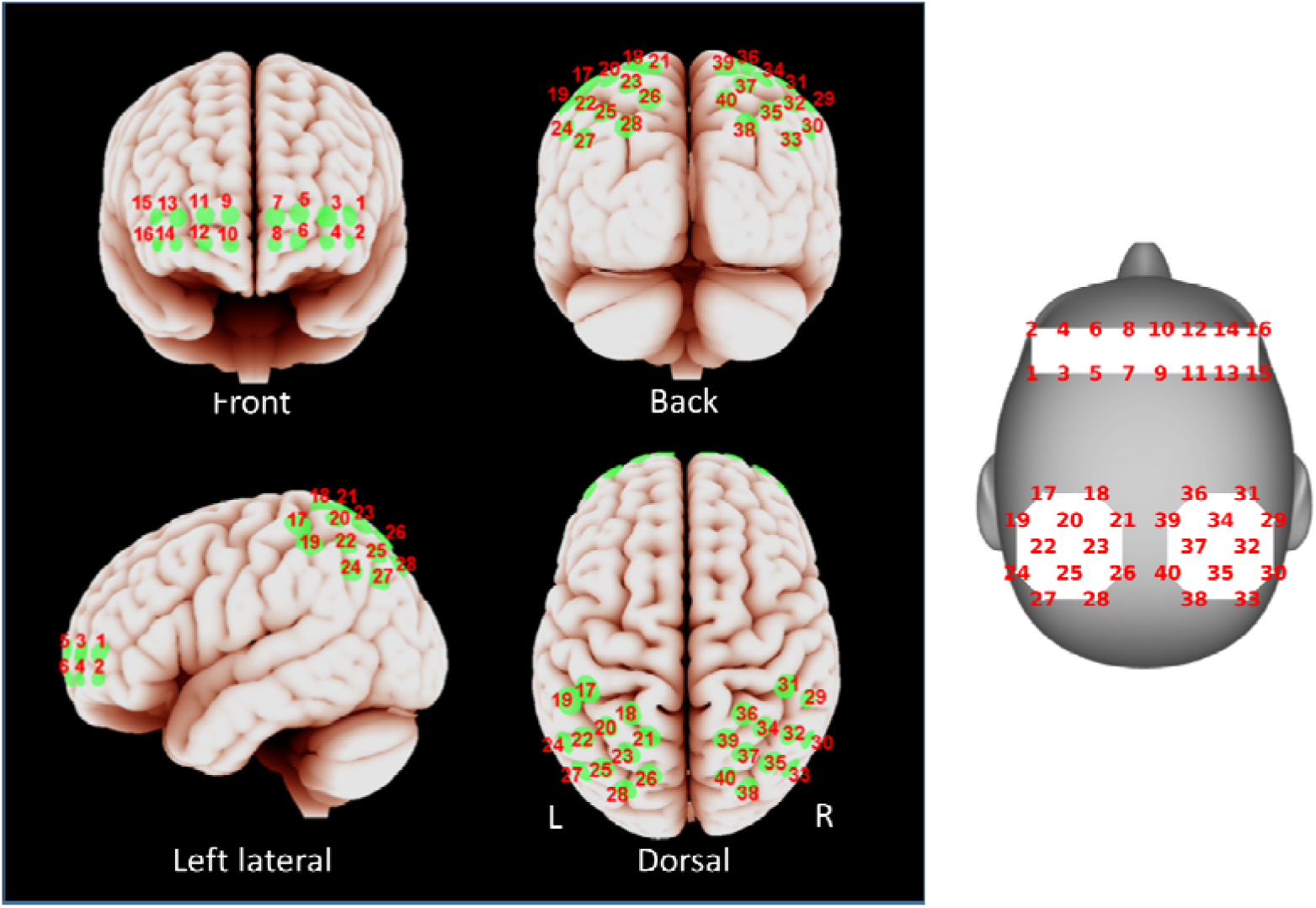
Optode locations. **Left**: Approximate spatial registration of optode locations to MNI space. **Right**: Schematic representation of the same optode locations on head surface which will be used later to show results. The virtual spatial registration approach was adopted to estimate the projection of the optodes on cortical surface in MNI space ^35^. The center of each green colored patch stands for the most likely position and the radius represents the standard deviation of displacement.

### fNIRS Data Preprocessing

fNIRS raw light intensity signals were converted to changes in oxygenated hemoglobin (ΔHbO) and nd deoxygenated hemoglobin (ΔHbR) concentrations using the modified Beer-Lambert law^39^. The raw signal and nd hemoglobin concentration changes were inspected both visually and also using the automated SMAR algorithm ^40^, which uses a coefficient-of-variation based approach to assess signal quality, reject problematic channels with bad contact or saturated raw light intensity. Next, the ΔHbO and ΔHbR time series for each optode and participant were band-pass filtered (0.01-0.5 Hz) and down-sampled to 2Hz. We considered only the period from 15 to 399 seconds, with respect to story start, in the signal time courses from each audio story. The first 15s were rejected to account for subjects’ initial period of adjustment to the listening comprehension task, and 399 s is the duration of the shortest story. Prior to subsequent analysis, the signal time courses were standardized optode-wise to have a mean of 0 and a standard deviation of 1.

### fNIRS Analysis

#### Inter-subject correlation

We first evaluated the reliability of the correlation between listeners’ brain activity using an inter-subject multilevel GLM similar to the one adopted by Stephens *et al.*^12^ for each of the four conditions: E1, E2, T1 and T2. We expected neural coupling between listeners to emerge only for the English story conditions E1 and E2, as none of the subjects understood Turkish.

At the individual subject level, a GLM with AR(1) (first-order autoregressive) error model was estimated using the average time course of the listeners as the independent variable and the time course of an individual listener as the dependent variable as follows:

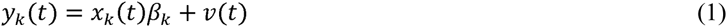

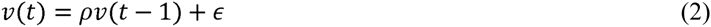

where *y*_*k*_(*t*) is the time course of a channel from listener k, *x*_*k*_(*t*) is the average time course of a channel from all the other listeners except for listener *k*, *ρ* is the autocorrelation coefficient and *ε~N*(0, *σ*^2^). AR(1) error model has been frequently adopted in event-related fNIRS analysis to model auto-correlated noise caused by low-frequency drift and physiological processes such as cardio-respiratory and blood pressure changes^41^.

At the group level, we tested the hypothesis *H*_1_:*β* > 0 using a one-tailed one-sample t-test evaluated on the slopes *βk* (*k* = 1, …, *n*) estimated at the individual level. We used the Benjamini–Hochberg procedure^22^ to control FDR among 80 statistical tests (2[ΔHbO/ΔHbR] × 40[optodes]) with *q* < 0.01.

#### Speaker-listener coupling

We evaluated the coupling between optode *i* of speaker and optode *j* of the listeners for all permutations of (*i,j*) (*N* = 40 × 40 *optodes* = 1600 *pairs*). The multilevel GLM was applied as it was for listener-listener coupling except that the time course of optode *i* of a speaker was used as independent variable *x*^*i*^(*t*) and the time course of optode j of each individual listener *k* was used as dependent variable 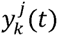.

#### Temporal asymmetry of coupling

The speaker-listener coupling analysis was repeated by shifting the speaker’s time course with respect to those of the listeners from −20s to 20s in 0.5s increments. At each time shift, FDR was controlled among 3200 statistical tests (2[ΔHbO/ΔHbR] × (40×40)[optode pairs]) with *q* < 0.01. As a comparison, listener-listener coupling was re-evaluated by shifting the average listener time series with respect to that of each individual listener.

### fMRI Analysis

The fMRI dataset included 17 subjects listening to story E2 (the “Pie-man” story), which was recorded and used in a previous study^14^. To compare with the fNIRS results, we considered only voxels from the outer layer of the cortex in the neighboring regions of prefrontal and parietal sites as shown in Figure 8. The “neighboring regions” are defined as voxels within a radius of 2.7 standard deviations from the center of each optode projection. A total of 994 voxels were chosen in this manner. The voxel time courses were high-pass filtered at 0.01 Hz (for comparison with fNIRS signals) and trimmed to only include 15-399 seconds (with respect to story start), and the inter-subject multilevel GLM described in section 0 was employed for model analysis. FDR was controlled among the 994 voxels with *q* < 0.01.

### fMRI-fNIRS Correlation

To estimate the correlation between BOLD and fNIRS signals, both signals were high-pass filtered at 0.01Hz. The BOLD time courses were z-scored normalized for each voxel and then averaged across the 17 subjects. fNIRS time courses were down-sampled to 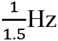 to match the fMRI sampling rate. The inter-subject multilevel GLM approach was then applied with the time course of optode *i* of an fNIRS subject *k* as independent variable 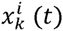 and the time course of voxel *j* averaged across fMRI subjects as dependent variable *y*^*j*^(*t*).

The aforementioned procedure was applied first to estimate all possible correlations between channels and their corresponding voxels in the left prefrontal (8 optodes × 131 voxels), right prefrontal (8 optodes × 144 voxels), left parietal (12 optodes × 376 voxels) and right parietal (12 optodes × 343 voxels) areas. The voxels were selected as described in section 0. To correct for multiple comparisons, FDR was controlled with a threshold of 0.05.

## Author Contributions

YL performed the experiment, collected the fNIRS data, analyzed the data and prepared/wrote the manuscript. EP analyzed the data and prepared/wrote the manuscript. ES collected the fMRI data, and participated in manuscript revision. PAS provided advise on the statistical approach and participated in manuscript revision. BO supported the idea and participated in manuscript revision. UH supported the idea, designed the experiment, discussed and interpreted the results and revised the manuscript. HA initiated and supervised the study, designed the experiment, analyzed the data, discussed and interpreted the results as well as prepared/revised the manuscript.

